# Evaluating Improvements to Exposure Estimates from Fate and Transport Models by Incorporating Environmental Sampling Effort and Landscape-level Contaminant Use

**DOI:** 10.1101/472969

**Authors:** Samantha L. Rumschlag, Scott M. Bessler, Jason R. Rohr

## Abstract

The Pesticide in Water Calculator (PWC) is a fate and transport model used by the Environmental Protection Agency and Health Canada to estimate pesticide exposures in lentic freshwater ecosystems and make pesticide registration decisions. We leverage over 600,000 field measurements of 31 common insecticides and herbicides to test whether incorporating environmental sampling effort (number of times a pesticide is sampled) and landscape-level contaminant use (national application amount) can improve PWC validation and prediction, respectively. We found that maximum measured concentrations of 38% of herbicides and 42% of insecticides exceeded maximum estimated environmental concentrations (EECs) generated by the PWC, suggesting that EECs often do not represent worst-case exposure. For lentic systems, variance in pesticide field measurements explained by EECs increased by 50% when sampling effort was included. For lotic systems, variance explained increased by only 4%, most likely because lotic systems are sampled over 4.9 times as much as lentic systems. Including landscape-level use more than doubled the ability of the PWC to predict maximum pesticides concentrations in lentic systems. Exposure characterization in risk assessment can be improved by including sampling effort in model validation and landscape-level use in predictions, thus providing more defensible environmental standards and regulations.

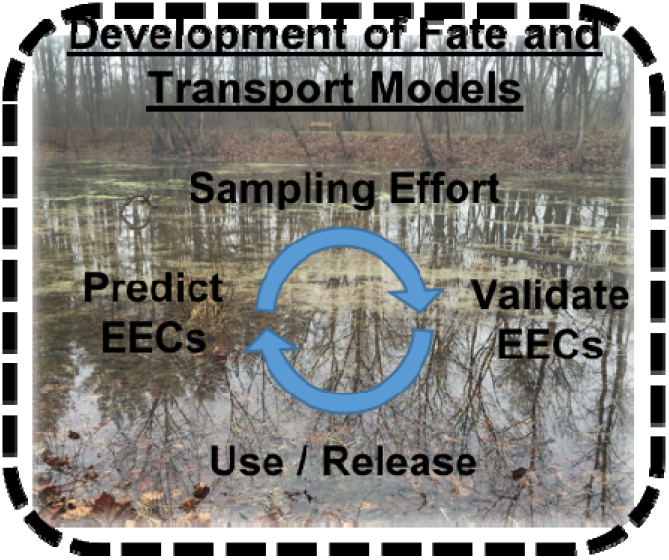

## INTRODUCTION

Chemical pollution represents one of the most widespread and destructive forms of human disturbance on earth^1–3^, threatening the health and wellbeing of humans^4–6^ and the environment^7–9^. For instance, more than 500 million pounds of active ingredients of pesticides are applied annually in the US^10^, leading to well-documented, widespread contamination of freshwater systems^11–13^ that provide habitat for about 10% of all described taxa on earth^14^. Therefore, the ability to predict levels of contamination in the field is critical to accurately assessing human and wildlife exposures and designing effective management strategies to minimize risks in sensitive systems.

Fate and transport models are important tools for predicting contaminant exposures. For instance, the United States (US) Environmental Protection Agency (EPA) and Health Canada use the Pesticide in Water Calculator (PWC) model to generate a peak estimated environmental concentration (EEC)^15^ of a focal pesticide in a standardized lentic waterbody that is a set distance from a site of application^16^. The model calculates an EEC based on inputs of pesticide traits (e.g. half-life and Koc), application amount and frequency (based on crop of interest), and soil and climatic characteristics (based on a region of interest)^16^. EECs of a variety of chemicals, including those that are not pesticides, are used in human health and ecological risk assessments. Generally, the EPA uses the PWC to predict pesticide EECs in ponds and reservoirs, which are used in ecological risk assessments and drinking water assessments, respectively^17^. Historically, the maximum EEC for pesticides has been regarded as a “worst-case” chemical exposure scenario in freshwater systems by the EPA^15^. In risk assessment, EECs are compared against toxicity values (e.g. LC50) to characterize the likelihood of toxicity at a given level of exposure^15,18^. Evaluation of EECs in this way informs the development of environmental standards, policies, guidelines, and regulations, as well as the registration of chemicals for legal use^18^.

While the PWC might be the best tool available currently for decision makers to estimate the potential for pesticide contamination in freshwater ecosystems, the increased availability of large-scale data on pesticide use and detection provides an opportunity for the accuracy of these exposure estimates to be evaluated and improved. The true or actual peak environmental concentration of any given pesticide is an extremely rare event in time and space, and thus validating model predictions of peak environmental concentrations necessitates copious field measurements, many more than are required to reliably predict mean environmental concentrations^19^. Fortunately, over the last three decades, federal agencies including the EPA and US Geological Survey (USGS) have compiled hundreds of thousands of field measurements of pesticides from lotic (streams and rivers) and lentic (ponds and reservoirs) freshwater ecosystems across the US. These publicly available data allow us to evaluate if EECs are indeed indicative of worst-case exposure scenarios and to determine the congruence between predicted EECs from the PWC and measured maximum concentrations of pesticides in the field.

The development of fate and transport models often progresses by repeating the following two steps: (1) a validation step, where predicted maximum EECs are correlated with measured or observed maximum environmental concentrations to determine their accuracy^20,21^, and (2) a prediction improvement step, where the model is modified to improve its fit to measured field concentrations (Fig. 1A). We use the term validation to mean the process of comparing model output to measurements^22^. These two steps are the foci of the current study.

**Figure 1.**
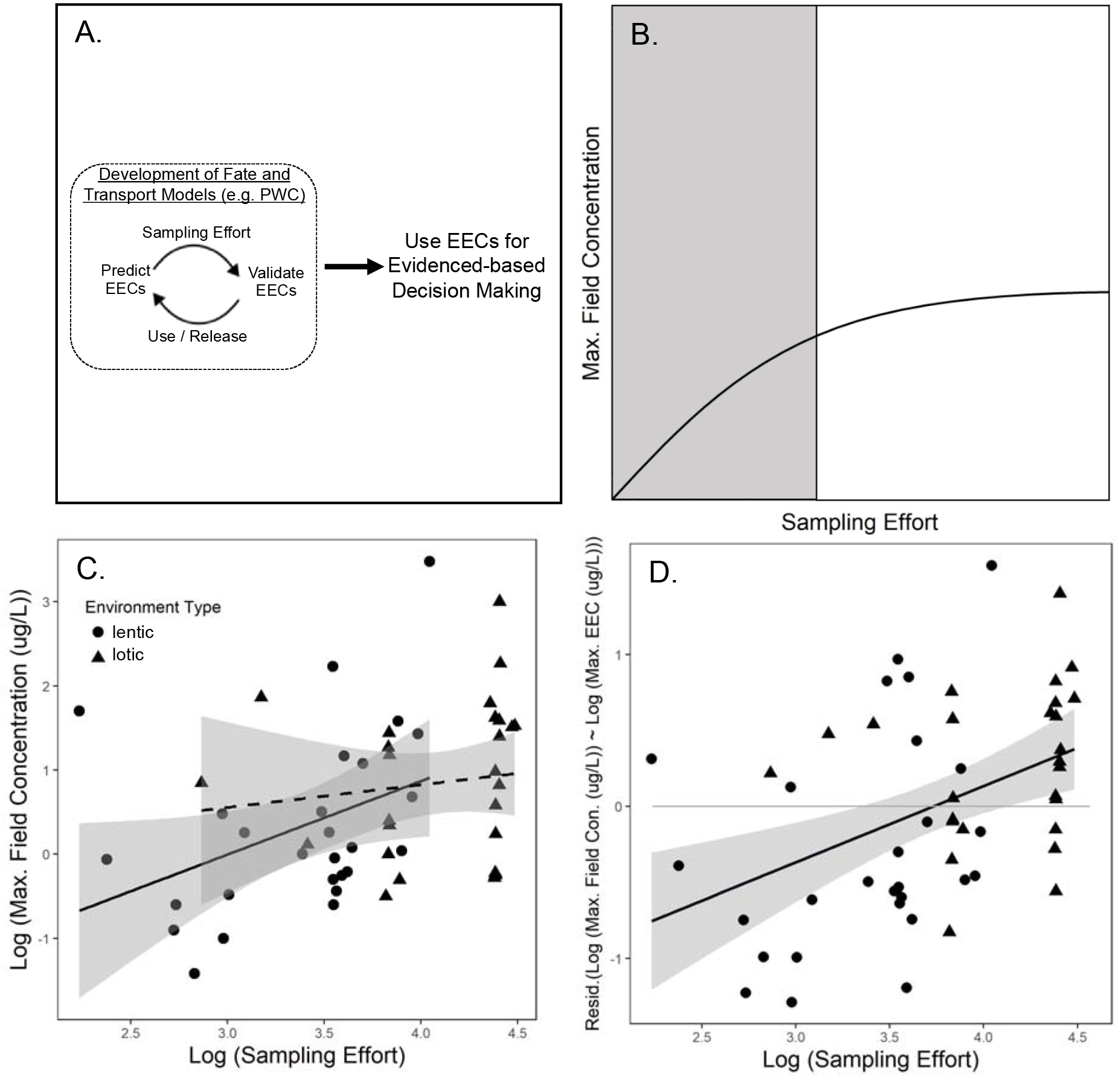
**A)** Conceptual model for improving fate and transport models like the Pesticide in Water Calculator (PWC), which produces estimated environmental concentrations (EEC) for pesticides. First, predicted EECs need to be validated using measured field concentrations to determine their accuracy. We predict that accounting for sampling effort will improve the fit between EECs and field measurements. Second, we will improve EECs by accounting for multiple sources of contaminant use or release. The accuracy of EECs is important because they used in federal decision making. **B)** Predicted asymptotic relationship between sampling effort and maximum (max.) field concentration. As sampling effort increases, the likelihood of detecting a peak concentration increases when sampling effort is at low to mid-levels as shown in gray. At mid to high levels of sampling effort, the influence of increased sampling effort on the likelihood of detecting a peak concentration reaches a limit, and no discernible relationship exists between sampling effort and max. field concentration as shown in white. We predict that sampling effort would account for more variance between maximum field concentration and maximum estimated environmental concentration (EEC) when sampling effort occurs in the lower range (in gray) compared to the higher range (in white). **C)** Observed relationship between sampling effort and maximum field concentration in lotic (circles, solid line) and lentic (triangles, dashed line) systems. Increased sampling effort is positively associated with maximum lentic concentration (Table 2, F = 4.552, *p* = 0.043) but not maximum lotic concentration (Table 2, F = 0.436, *p* = 0.515). The positive relationship for lentic systems matches the positive relationship at low to mid-sampling effort shown in gray in Figure 1B. The absence of a relationship for lotic systems matches the asymptote at mid to high sampling effort in Figure 1B. **D)** Observed relationship between sampling effort and the residuals of maximum field concentrations in lotic (circles) and lentic (triangles) in systems and EEC (Table 2, χ^2^ = 12.339, *p* <0.001). As sampling effort increases, the likelihood of a field concentration exceeding an EEC increases, which is represented by a positive residual. Gray band represent a 95% confidence interval, and a light gray reference line at 0 represents where maximum field concentration would equal maximum EEC.

An important consideration for model validation of contaminant fate and transport models might be environmental sampling effort, defined as the total number of times a pesticide is surveyed across locations and time. If we are to determine how well maximum EECs predict maximum field concentrations of contaminants, we must account for the variance in maximum field concentrations that is a function of sampling effort. For instance, we propose that sampling effort should be related asymptotically to the maximum environmental concentration of a contaminant, such that increases in sampling effort increase the likelihood of detecting the true peak environmental concentration at low (gray section in Fig. 1B) but not high sampling efforts (white section in Fig. 1B). Given this proposed relationship, we hypothesize that incorporating information on environmental sampling effort will improve the ability of the PWC to predict maximum environmental concentrations, but only for systems that are not well sampled and thus fall on the section of the sampling effort-maximum field concentration curve that is increasing rather than near the asymptote.

PWC predictions might be improved by accounting for multiple sources of contaminant use or release. Most fate and transport models, including the PWC, assume a single point source of contamination, but measured concentrations in freshwater ecosystems are often the result of runoff and aerial deposition from multiple sources of contamination across the landscape. The inaccurate assumption of a single point source could misrepresent the true or actual peak concentration in the environment that the PWC seeks to model. Thus, we hypothesize that incorporating information on landscape-level use or releases of chemicals might improve the ability of fate and transport models to predict maximum field concentrations because landscape-level use is a proxy for multiple sources of contamination. The USGS recently provided pesticide use estimates for each county in the US, allowing us to evaluate how the inclusion of landscape-level pesticide applications affects the ability of the PWC to predict maximum measured concentrations in the environment.

To test the hypotheses that the validation and predictions of fate and transport models can be improved by accounting for environmental sampling effort and landscape-level contaminant release information, respectively, we selected 31 of the most commonly used pesticides and compiled data describing their use, application rate, environmental mobility, EECs from the PWC, and maximum measured environmental concentrations in lentic and lotic systems. We predicted that EECs would not represent worst-case scenarios of exposure because EECs fail to incorporate landscape-level pesticide use and instead model a commonly unrealistic single point-source. Given the postulated importance of sampling effort, we predicted that the PWC would more accurately predict maximum concentrations in lotic than lentic systems because lotic systems are sampled for pesticides nearly 4.9 times as much as lentic systems (mean number ± standard deviation of lotic versus lentic samples per pesticide from federal databases: 16,111± 10,301 vs. 3,304 ± 3,005). Finally, we predicted that the PWC’s predictions of maximum EECs could be improved by incorporating landscape-level use or release information to account for likely multiple sources of pesticides to freshwater ecosystems.

## METHODS

### Pesticide Selection

Our analyses focus on the 31 most commonly used herbicides and insecticides applied on corn in the US (Table 1). To select this group of pesticides, we first ranked insecticides and herbicides based on their estimated use in the US by summing 2006 county-level pesticide use estimates from the Estimated Annual Agricultural Pesticide Use dataset provided by Pesticide National Synthesis Project of the National Water Quality Assessment (NAWQA) Program (US Geological Survey [USGS]) (https://water.usgs.gov/nawqa/pnsp/usage/maps/county-level/). We classified each pesticide as an herbicide or insecticide using the primary use type classifications indicated by the Pesticide Action Network (PAN) Pesticide Database (http://www.pesticideinfo.org/). We excluded mineral or biologic (e.g. bacteria) pesticides, because we were interested in examining the transport and fate of synthetic compounds. From these most commonly used synthetic herbicides and insecticides, we selected compounds that were detected in streams from 1992 to 2012 by the USGS NAWQA program (www.waterqualitydata.us/portal, obtained on 30 March 2017). Finally, we examined commercial product use labels and only included compounds that were used on corn because standard EPA scenarios used in the calculation of EECs (see below) are more frequently available across geographic regions in the US for corn than other crops. This selection process resulted in 16 herbicides and 15 insecticides (Table 1).

**Table 1.** List of pesticide active ingredients and type included in the present analyses. Pesticide abbreviations are used as point labels in the subsequent figures.

| Pesticide Active Ingredient | Pesticide Abbreviation | Pesticide Type |
| --- | --- | --- |
| 2,4-D | 24D | herbicide |
| Acetochlor | ACE | herbicide |
| Alachlor | ALA | herbicide |
| Atrazine | ATR | herbicide |
| Bromoxynil | BRO | herbicide |
| Dicamba | DIC | herbicide |
| Dimethenamid | DID | herbicide |
| Diuron | DIU | herbicide |
| Glyphosate | GLY | herbicide |
| MCPA | MCP | herbicide |
| Metolachlor | MET | herbicide |
| Metribuzin | MTR | herbicide |
| Oxyfluorfen | OXY | herbicide |
| Pendimethalin | PEN | herbicide |
| Simazine | SIM | herbicide |
| Trifluralin | TRI | herbicide |
| Aldicarb | ALD | insecticide |
| Carbaryl | CAR | insecticide |
| Carbofuran | CBO | insecticide |
| Chlorpyrifos | CHL | insecticide |
| Clothianidin | CLO | insecticide |
| Diazinon | DIA | insecticide |
| Dimethoate | DIM | insecticide |
| Imidacloprid | IMD | insecticide |
| Malathion | MAL | insecticide |
| Methomyl | MTH | insecticide |
| Methyl Parathion | MLP | insecticide |
| Phorate | PHO | insecticide |
| Propargite | PRO | insecticide |
| Tefluthrin | TEF | insecticide |
| Terbufos | TER | insecticide |

### Building a Dataset Characterizing Herbicides and Insecticides

We built a dataset describing each selected pesticides’ use, application rate, environmental mobility and persistence, and maximum measured environmental concentration (Tables S1 and S2). For each compound, we determined an estimate of national use by summing all county-level pesticide estimates from the Estimated Annual Agricultural Pesticide Use dataset from 1992 to 2012. Maximum concentrations of pesticides in lotic systems were taken from stream survey data from 1992 to 2012 from the USGS NAWQA program (from https://www.waterqualitydata.us/, obtained on 30 March 2017, filtered by NAWQA program and stream site type). The total number of stream surveys from which maximum concentrations were taken totaled 499,435. Maximum concentrations of pesticides in lentic systems were taken from surveys of lakes, reservoirs, impoundments, and wetlands from 1992 to 2012 available from National Water Quality Monitoring Council (https://www.waterqualitydata.us/, obtained on 9 November 2017, filtered by site type to include lakes, reservoirs, impoundments, and wetlands). The total number of surveys from these lentic systems from which maximum concentrations were taken totaled 129,471. Although a valuable consideration might be to examine a distribution of estimated environmental or field concentrations and focus on the top 95% or 99% percentile, risk assessments are generally concerned with a single maximum estimated environmental or field concentration, so our focus was on gathering a single maximum concentration for each pesticide. For each pesticide, a single maximum concentration was taken from across lentic and lotic survey locations and times. To help limit the influence of timing of sampling on detection of maximum concentrations, we excluded samples that were triggered by a hydrologic event (i.e., event-based sampling), such as a flood or a storm. Instead, we focused on field samples that were gathered as part of routine-based sampling efforts. Since we wanted to record maximum observed pesticide concentrations; both filtered and whole water sample were considered. We also recorded sampling effort for each pesticide in lentic and lotic systems, which was the number of times a pesticide was surveyed for across locations and time. More information concerning how each maximum pesticide concentration was determined is provided in Tables S3 and S4. In addition, we gathered maximum field concentrations from lakes, ponds, agricultural ditches, and tailwaters by reviewing the published scientific literature to evaluate whether maximum EECs are indeed worst-case scenarios of exposure using the most information possible on maximum lentic concentrations. We conducted a literature search using Web of Science and Google Scholar using combinations of the following terms: “concentration”, “tailwater”, “pond”, “ditch”, “runoff”, “field concentration”, and the name of the focal pesticide (e.g. atrazine). In the final dataset, we include only values from the literature that exceeded pesticide field database values in lentic systems. Individual maximum concentrations of pesticides gathered from databases or the literature represent observed maximum measured concentrations and not the true or actual peak concentrations, which can only be greater than or equal to the maximum measured concentration^19^.

### Generating Estimated Environmental Concentrations

Data describing the environmental mobility and persistence of herbicides used in the calculation of EECs, including Koc, water column metabolism half-life, benthic metabolism half-life, foliar half-life, aqueous photolysis half-life, molecular weight, vapor pressure, and solubility, were taken primarily from the Pesticide Properties DataBase from the University of Herfordshire (PPDB, https://sitem.herts.ac.uk/aeru/ppdb/en/). Values for hydrolysis half-life and aerobic soil half-life were taken from PAN Pesticide Database. When values were not available for certain pesticides from PAN or PPDB, we used data from the Toxicology Data Network (TOXNET) from the National Institutes of Health (https://toxnet.nlm.nih.gov/newtoxnet/hsdb.htm) as indicated in Tables S1 and S2.

Additional pesticides traits included Henry’s constant, heat of Henry, air diffusion coefficient, and application information (Tables S1 and S2). Henry’s constant and the heat of Henry were taken from the EPA’s Estimation Program Interface (EPI) Suite, specifically HENRYWIN. Henry’s constant was calculated using the bond contribution method. We calculated the air diffusion coefficient using the EPA’s On-line Tools for Site Assessment Calculation (https://www3.epa.gov/ceampubl/learn2model/part-two/onsite/estdiffusion-ext.html). Data concerning number of applications per year, timing of applications, and maximum recommended application rate and method were taken from US commercial pesticide product labels. For herbicides, product instructions for pre-emergent applications for corn were followed when available. We assumed that the last application of pre-emergent herbicides would occur just after planting, 12 days prior to corn emergence. For herbicides that are exclusively applied post-emergence, we assumed applications would occur 10 days after corn emergence. We assumed all herbicides would be applied by direct ground spray, unless product labels indicated the need for soil incorporation. In those cases, applications were set to occur at the suggested depth of soil incorporation based on the product label. For insecticides, product application instructions for post-emergent applications for corn were used when available. We assumed that the first applications would occur 30 days after emergence by spray above the plant. For insecticides that are applied pre-emergence, we assumed applications would occur 12 days before emergence by ground spray at the depth of soil incorporation according to the product labels.

Using the EPA’s Pesticide in Water Calculator v. 1.52 (PWC, https://www.epa.gov/pesticide-science-and-assessing-pesticide-risks/models-pesticide-risk-assessment#PWC), we generated EECs of the selected pesticides. Model inputs consisted of mobility, persistence, and application data for individual pesticide compounds (Tables S1 and S2). For all pesticide compounds, water, benthic, and soil reference temperatures were assumed to be 23 degrees C, and photolysis reference latitude was 40 degrees. When foliar half-life was not available for a given pesticide, foliar half-life was assumed not to be a large contributor to breakdown in the environment in the PWC model and was set to zero. Under the recommendation of the PWC user manual, efficiency was set to 0.99 and drift was set to 0.01 for all pesticide compounds. Applications were assumed to occur every year. For each pesticide compound, EECs were generated for both ponds and reservoirs in each of five different states (Illinois, Mississippi, North Carolina, Ohio, and Pennsylvania), which varied in their meteorological and geological model inputs provided by the PWC software. This resulted in 10 EECs values for each pesticide. We used the maximum EEC of these 10 estimates for each pesticide in all statistical analyses.

### Statistical Analyses

To determine how often maximum EECs represent worst-case scenarios of pesticides in lentic systems, we calculated the proportion of pesticides for which the maximum environmental concentrations in lentic systems exceeded maximum EECs from PWC models. In this evaluation, the point of comparison for the EEC was the highest concentration of pesticide found in the National Water Quality Monitoring Council database or in the literature. We incorporated literature and database field measurements because we wanted to use all possible available data to describe maximum lentic field values. In all other analyses, we use maximum lentic field values from the National Water Quality Monitoring Council exclusively to ensure that the methods of estimating maximum lentic and lotic field concentrations were similar, which is an important consideration for the quantitative assessment for model validation and improvement of model predictions. The literature concentrations had to be removed from these analyses because they did not use consistent sampling methodology across studies.

To evaluate the effects of sampling effort on detection of maximum field concentrations in lentic and lotic systems, we built two separate linear models (lm function, *stats* package^23^) in which the response was either maximum lentic or lotic concentration and the predictor was sampling effort, defined as the total number of times a pesticide was surveyed for between 1992 and 2012 respective to each system, including surveys which resulted in no detection of the pesticide. To evaluate if inclusion of sampling effort improved model validation of maximum EECs with maximum field concentrations, first we examined the effect of sampling effort on the relationship between maximum field concentration and maximum EEC. We extracted the residuals from a mixed model (lmer function, *lme4* package^24^) with maximum field concentration as the response and maximum EEC as the predictor with pesticide compound as the random effect. These residuals became the response in a subsequent mixed model, where the predictor was sampling effort, and the random effect was pesticide compound. Next, we compared models predicting maximum field concentrations from maximum EECs with and without observations weighted by sampling effort. We constructed linear models (lm function, *stats* package^23^) in which the response was either maximum field concentration detected in lentic (from NAQWA) or lotic systems (from National Water Quality Monitoring Council) and the predictors were maximum EEC, pesticide type (insecticide or herbicide), and the interaction between these two predictors. We ran each model with and without weighting observations by sampling effort. In the evaluation of the effect of maximum field concentration on maximum EEC in this set of analyses, we used a one-tailed hypothesis test because of the prediction that maximum field concentration would be positively associated with maximum EEC. To compare the amount of variance explained by each model, we calculated adjusted-*R^2^* values.

Lastly, we sought to evaluate if the ability of EECs to predict field concentrations in lentic systems could be improved by including landscape-level pesticide use and release as a predictor. We focus on improving EECs in reference to lentic field concentrations because the EPA uses the PWC to predict pesticide EECs in ponds and reservoirs for ecological and drinking water risk assessments, respectively^17^. We used multimodel inference (*MuMIn* package^25^, which fits models using combinations of all predictors given in a global model and ranks candidate models by second-order Akaike Information Criteria corrected for small sample sizes (AICc) (dredge function). In our global model, the response was maximum lentic concentration (from the National Water Quality Monitoring Council) and the predictors included: maximum EEC, pesticide type, pesticide use, all two-way and three-way interactions between these factors. Since our purpose was to improve the ability of EECs to predict field concentrations, we only considered candidate models that included maximum EEC as a predictor. To compare the influence of model factors across all candidate models, Akaike weights for each factor were summed across models to determine relative importance scores^26^. To evaluate the amount of variance explained by the top model, we calculated adjusted-*R^2^* values.

In all statistical models in the present analyses, all continuous variables were log_10_-transformed to meet assumptions of the analyses. The data analyzed contained the 27 pesticides found in lentic systems when analyses pertained exclusively to lentic data or when lentic and lotic data were combined. Analyses of all 31 pesticides occurred when lotic data were examined exclusively (e.g. for evaluation of inclusion of sampling weights for model validation of EECs with lentic field concentrations). For all models to determine if the predictors significantly influenced the responses, we used the Anova function in the *car* package^27^ (α=0.05). Figures were generated using *visreg*^28^ and *ggplot*^29^ packages. *R* 3.2.1 statistical software^23^ was used for all analyses.

## RESULTS

### Do EECs represent worst-case scenarios of pesticides in lentic systems?

Historically, EECs have been described as worst-case environmental concentrations^15^. However, maximum concentrations in lentic systems exceeded EECs for 37.5% of herbicides (6 of 16) and 41.7% of insecticides (5 of 12), suggesting that for many pesticides, EECs did not represent worst-case scenarios of exposure in lentic systems.

### What is the effect of sampling effort on detection of maximum field concentrations?

We hypothesized that maximum field concentration would increase asymptotically with sampling effort (Fig. 1B). As sampling effort increases, detected maximum field concentration should increase up to a point (gray section of Fig. 1B), after which increased sampling effort should have little to no association with maximum field concentration (white section of Fig. 1B). We observed this dichotomy in sampling effort according to environmental systems. Sampling effort was positively associated with maximum field concentration in lentic but not lotic systems (Fig. 1C, Table 2), most likely because sampling effort for pesticides in lentic systems represents a lower range of values compared to sampling effort in lotic systems. Lotic systems were sampled 4.9 times as much as lentic systems (mean number ± standard deviation of lotic versus lentic samples per pesticide: 16,111± 10,301 vs. 3,304 ± 3,005). Thus, observations from lentic systems seem to fall on the section of the hypothesized curve with a positive slope where increased sampling is associated with higher detected maximum field concentrations (i.e. gray section of Fig. 1B). In contrast, observations from lotic systems seem to fall on the section of the curve closer to the asymptote, so increases in sampling effort only have marginal effects on the maximum field concentration (i.e. white section of Fig. 1B). Following this pattern, we predicted that including sampling effort would improve model validation for maximum EECs in lentic but not lotic systems.

**Table 2.**
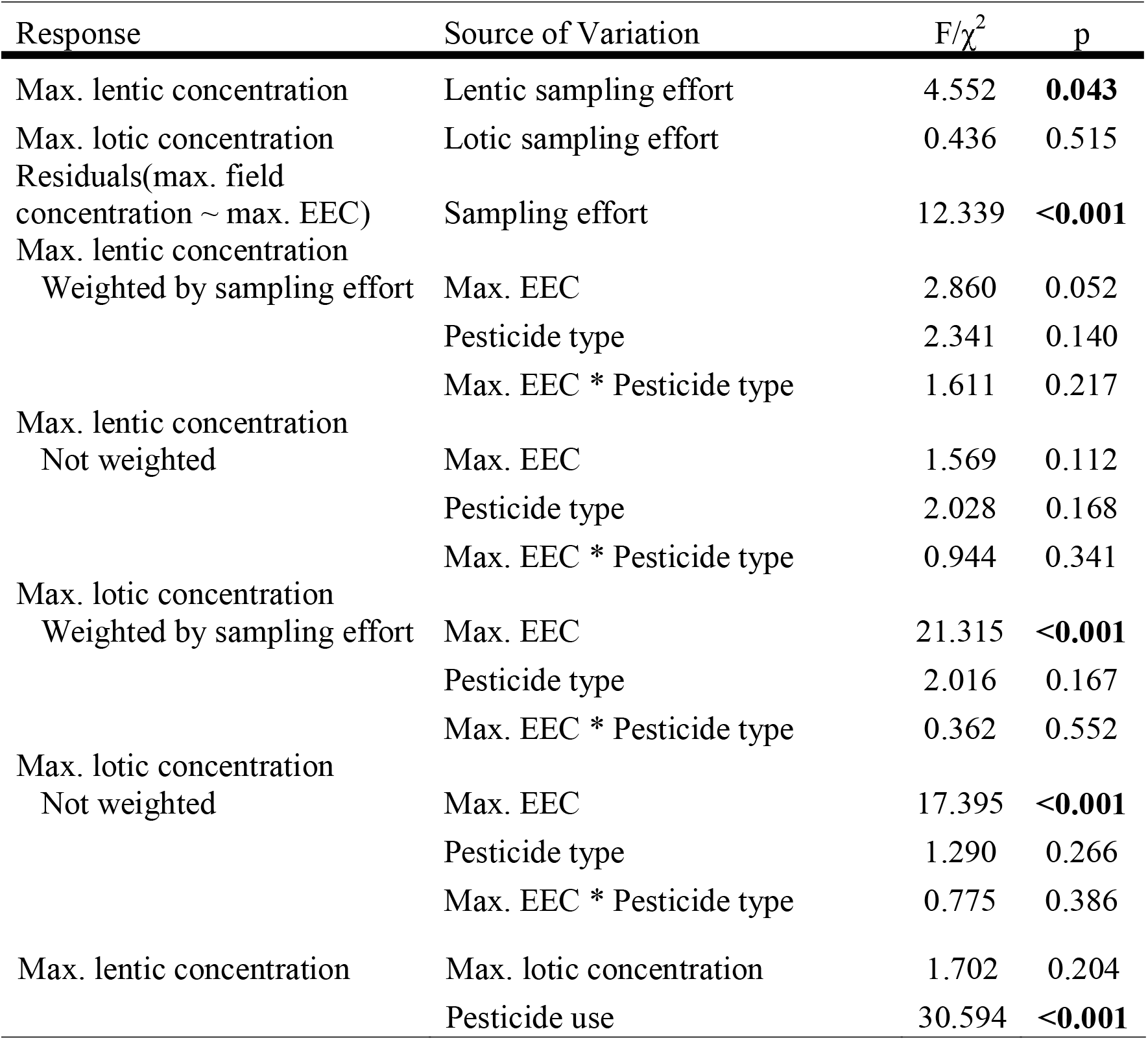
Analyses summaries examining 1) the influence of sampling effort on maximum (max.) lentic concentration, lotic concentration, and the residuals of maximum field concentration predicted by maximum estimated environmental concentration (EEC), and 2) the influence of maximum EECs on maximum lentic and lotic concentrations with and without sampling effort weighted. In this set of analyses, we used one-tailed tests for the effect of max. EEC on field concentrations. Finally, 3) we include a summary of the best fitting model predicting maximum lentic concentrations from model selection. *P*-values less than 0.05 are bolded. χ^2^ statistics correspond with a mixed model. F statistics correspond with non-mixed models. The data analyzed contained the 27 pesticides detected in lentic systems for all analyses, excluding evaluations between maximum lotic concentration and maximum EEC that included all 31 pesticides.

### Can inclusion of sampling effort improve model validation of maximum EECs with maximum field concentrations?

These differences in the association between sampling effort and maximum field concentration lead us to test if inclusion of sampling effort could improve model validation of maximum EECs with maximum field concentrations. In other words, we wanted to evaluate if incorporating sampling effort into models increases the variance in maximum field concentrations that can be explained by maximum EECs. First, we examined the influence of sampling effort on the relationship between maximum field concentration and EECs. We observed a positive effect of sampling effort on the residuals of a model predicting maximum field concentrations from maximum EECs (Fig. 1D, Table 2). At low to medium relative levels of sampling effort (log_10_ (sampling effort) = 2.24 to 3.78), maximum EECs tend to overestimate observed maximum field concentrations, which is represented by negative residuals, and at medium to high relative levels of sampling effort (log_10_ (sampling effort) = 3.78 to 4.57), maximum EECs more often underestimate maximum field concentrations, which is represented by positive residuals (Fig. 1D).

Next, we sought to evaluate if the inclusion of sampling effort could increase the amount of variance explained in maximum field concentrations from lentic and lotic systems by maximum EECs, an important consideration in validation of EECs. As hypothesized, sampling effort improved the fit of maximum EECs to maximum field concentrations for lentic systems more so than for lotic systems (Fig. 2, Table 2). The maximum EECs from the PWC, which are purported to represent maximum concentrations of pesticides in ponds and reservoirs, were not a significant predictor of maximum measured pesticide concentrations in lentic systems without weights but became nearly significant when weighting by sampling effort (Table 2). In fact, weighting observations by lentic sampling effort increased the relative amount of variance explained by 50% (Fig. 2A [Adjusted *R^2^* = 0.27], Fig. 2B [Adjusted *R^2^* = 0.18]). For lentic models with and without sampling effort weighted, while there was a positive trend between herbicide EECs and measured concentrations of herbicides in lentic systems, there was no discernible relationship between insecticide EECs and lentic insecticide concentrations (Fig. 2A, B). In other words, maximum EECs were a poor predictor of field concentrations for insecticides in lentic systems. For lotic systems, weighting observations by sampling effort increased the relative amount of variance explained by only 4% (Fig. 2C [Adjusted *R^2^* = 0.54], Fig. 2D [Adjusted *R^2^* = 0.52]). Maximum EECs were a significant positive predictor of maximum measured concentration of herbicides and insecticides in lotic systems regardless of whether we weighted by sampling effort or not (Table 2, Fig. 2C,D).

**Figure 2.**
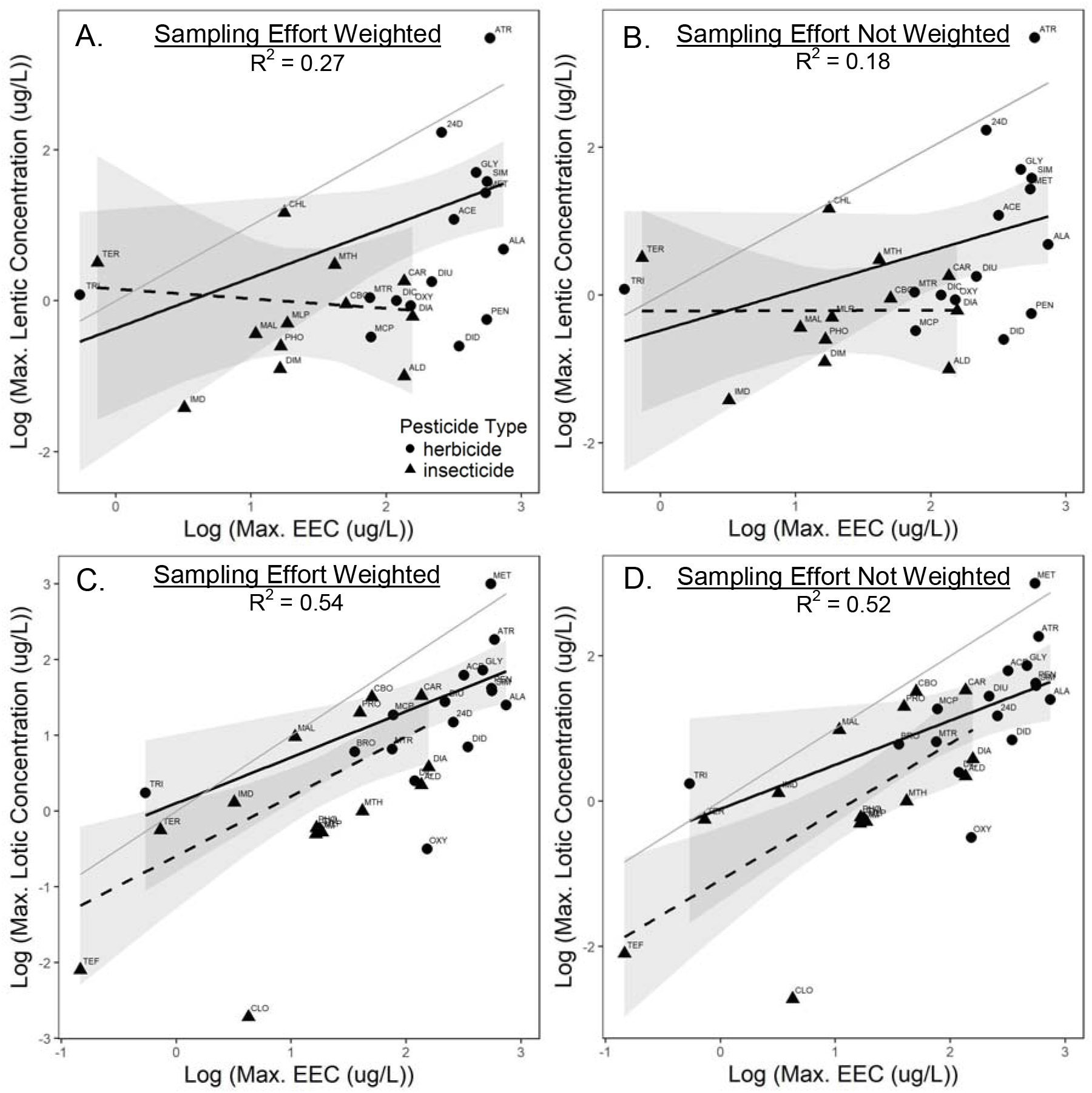
Associations between herbicide and insecticide maximum (max.) estimated environmental concentrations (EEC) and measured maximum field concentrations in lentic (A and B) and lotic (C and D) systems. Models were built with (A and C) and without (B and D) observations weighted by sampling effort. The association between maximum EEC and maximum field concentration is significant for the lotic system with and without observations weighted by sampling effort (Table 2, *p* <0.001, C and D) and nearly significant for lentic system when observations are weighted by sampling effort (Table 2, *p*=0.052, A and B). In all panels herbicides are shown with solid circles and solid lines, and insecticides are shown with triangles and dashed lines. Individual pesticides are labeled above and to the left of the point (see Table 1 for abbreviations). Gray bands represent 95% confidence intervals, and light gray lines are 1:1 references lines.

### Can EEC predictions be improved by including landscape-level pesticide use and release?

To test the hypothesis that inclusion of landscape-level contaminant use and release could improve the ability of maximum EECs to predict maximum field concentrations, we used model comparison techniques. Based on model comparison, the best-fitting model of maximum measured concentrations of pesticides in lentic systems included maximum EEC and estimated national use (model weight = 0.42). In this best-fitting model, estimated national pesticide use but not maximum EEC significantly predicted maximum measured concentrations of pesticides in lentic systems (Table 2). In addition, maximum EEC and estimated national pesticide use had the greatest relative importance scores (Fig. 3A). This best-fitting model more than doubled the ability of the PWC to predict maximum concentrations of pesticides in lentic systems (Adjusted *R^2^* = 0.64 vs. Adjusted *R^2^* = 0.27). Estimated national pesticide use was positively associated with maximum lentic concentration suggesting that pesticide use improves EEC predictions of herbicides and insecticides (Fig. 3B).

**Figure 3.**
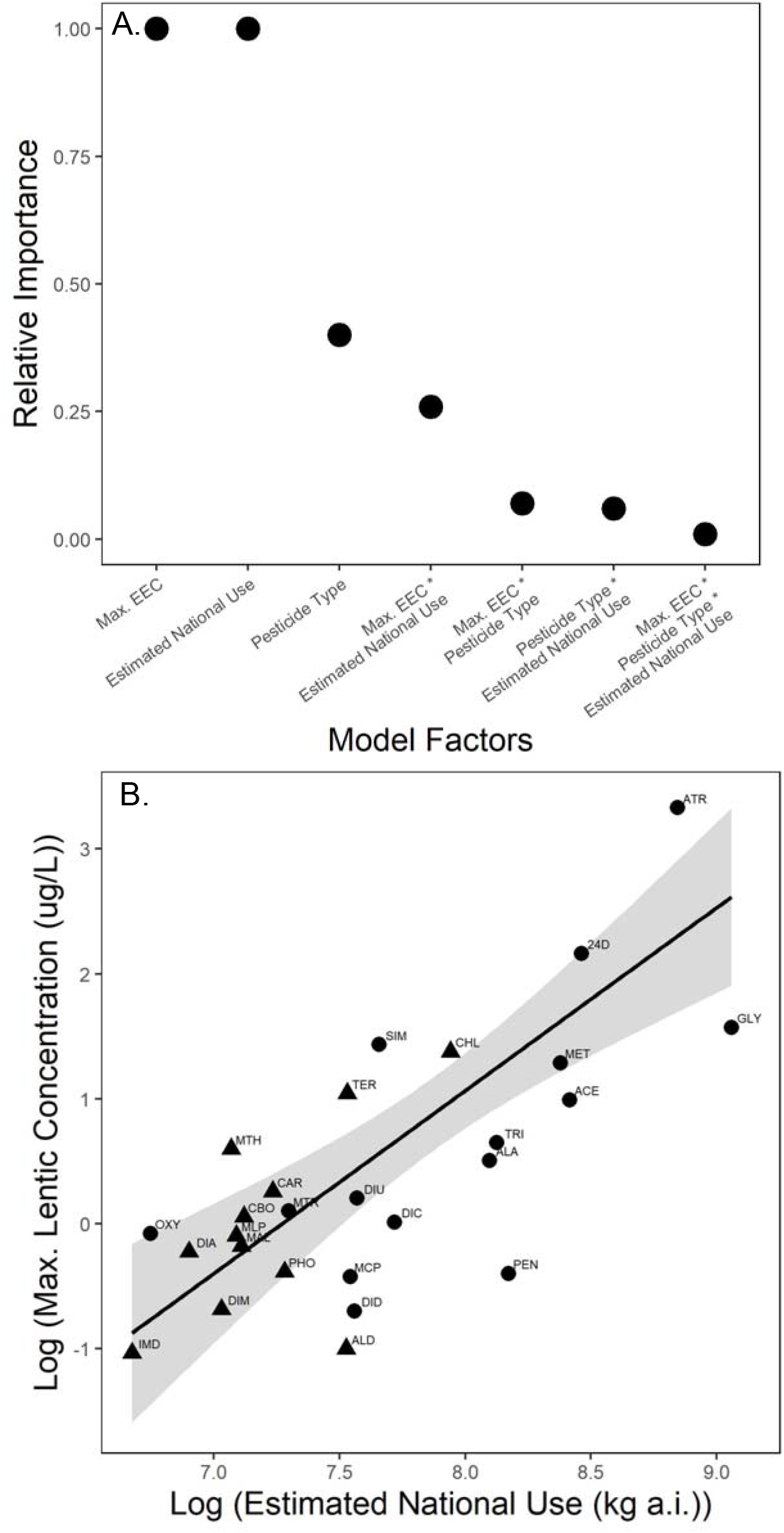
**A)** Relative importance scores of factors from model comparisons, evaluating the best predictors of maximum concentration of pesticides in lentic systems. Maximum estimated environmental concentration is abbreviated as Max. EEC. **B)** Conditional plot displaying the significant effect of estimated national pesticide use on maximum (max.) lentic concentration, controlling for maximum EEC, soil half-life, and pesticide type (based on best fitting model, Table 2, F = 30.594, *p* <0.001). Gray bands represent 95% confidence intervals. Conditional plot was generated using the *visreg* package in *R*.

## DISCUSSION

From an ecological risk assessment perspective, the ability to accurately predict concentrations of chemical contaminants is essential for the creation of defensible environmental standards, policies, guidelines, and regulations^18^. By leveraging over 600,000 field measurements of the most commonly used insecticides and herbicides, we use the PWC model as a case study to evaluate how to improve contaminant fate and transport models more generally. Consistent with our hypotheses, we demonstrate that incorporating environmental sampling effort and landscape-level contaminant use or release improves model validation and prediction, respectively, an approach that can be applied to other fate and transport models. Inclusion of sampling effort in model validation greatly improves the ability of EECs to predict the variance of field concentrations in poorly sampled lentic systems but only marginally improves prediction in well-sampled lotic systems. In addition, inclusion of landscape-level pesticide use as a measurement of multiple contaminant point-sources more than doubles the ability of the PWC model to predict maximum concentrations of pesticides in lentic systems.

### Model Validation: The Importance of Sampling Effort on the Ability of PWC Models to Predict Field Concentrations

When compared against maximum lentic field measurements, maximum pesticide EECs produced by PWC models for ponds and reservoirs perform poorly. For instance, historically, maximum EECs have been considered worst-case scenarios of exposure^15^, but our results show that this is a mischaracterization. If a maximum EEC is truly a worst-case scenario of exposure, we would expect that field concentrations of pesticides would never fall above an EEC, but for about ~40% of the most commonly used pesticides measured, field values exceed EECs. This finding is important because if risk assessors and policy makers consider maximum EECs as worst-case concentrations to gauge the greatest potential for toxicity, they would be underestimating levels of field exposures in many cases. This difference between maximum EECs and maximum field measurements indicates the need for improved model validation and prediction.

Testing the ability of EECs to predict field concentrations is an important step of model validation and model development^20,21^. Patterns of the observed relationship between sampling effort and maximum detected field concentrations lead us to the hypothesis that the importance of sampling effort on the ability of EECs to predict field concentration likely varies with lotic versus lentic systems because of differences in the amount of pesticide sampling effort in each system. For instance, lentic systems are sampled about 4.9 times as much as lotic systems. Because the relationship between sampling effort and maximum field concentration in lotic systems is positive, we hypothesized that sampling effort would be important for EECs to predict field concentrations in this system. In contrast, because sampling effort only has marginal effects on maximum field concentrations in lotic systems, we predicted that sampling effort would have little to no effect on the ability of EECs to predict field concentrations.

Consistent with our hypothesis, we show that the ability of maximum EECs to predict maximum field concentrations can be improved by weighting observations by sampling effort in both lentic and lotic systems, but the magnitude of this improvement is greater for lentic than lotic systems. Weighting observation by sampling effort increased the relative amount of variance explained by 4% for lotic systems and 50% for lentic systems. Consequently, these results demonstrate that accounting for contaminant sampling effort is an important component of model validation, especially when sampling efforts fall within the range in which sampling effort is positively corelated with maximum field concentrations. If scientists validate EECs by comparing maximum EECs to maximum environmental concentrations in order to determine if EECs are accurate or not, they must account for the variance in maximum environmental concentrations that are a function of sampling effort. By accounting for sampling effort, scientists can more accurately determine if EECs are valid approximations of contaminant exposures.

For insecticides in lentic systems, even though the variance explained in maximum field concentrations by maximum EEC increases when we accounted for sampling effort (as represented by a shift in the dotted line closer to the 1:1 reference line in Fig. A compared to Fig. B), the ability of EECs to predict field concentrations was still poor (shallow slope of the dotted lines in Fig. A. and B). The inability of the maximum EECs to predict maximum field concentrations of insecticides compared to herbicides might be a function of pesticide use. Use of herbicides is about five times greater than insecticides in the US^30^, and so the power to detect an association between maximum herbicide EECs and maximum herbicide field concentrations should be greater than that for insecticides. As a result, maximum field concentration of herbicides might be closer to the true peak concentrations compared to insecticides.

### Improving EEC Predictions with Landscape-level Use and Release

Even when field concentrations are the result of intensive sampling, maximum EECs can still underestimate maximum field concentrations (which is represented by positive residuals in Fig. 1D). The assumption of a single point source likely results in this underestimation of the peak environmental concentrations by EECs. For instance, most fate and transport models, including the PWC, assume a single point source of contamination, but measured concentrations of contaminants in freshwater ecosystems are often the result of runoff and aerial deposition from multiple sources of contamination across the landscape.

With this motivation, we attempted to improve the ability of EECs to predict field concentrations in lentic systems by accounting for landscape-level pesticide use. For both herbicides and insecticides, landscape-level pesticide use improved the ability of EECs to predict maximum concentrations in lentic systems, more than doubling the variance explained compared to a model without landscape-level use. Most notably, when the model accounted for sampling effort and pesticide use, the ability of EECs to predict maximum field concentrations in lentic systems went from no relationship (Fig. 2A) to a significant positive relationship (Fig. 3B). Improvement in EECs by inclusion of pesticide use is what we would predict if environmental pollution is the result of multiple point sources of contamination. These results suggest that pesticide use at the national level is likely an improved indicator of pesticide loading into a freshwater ecosystem than the single point-source of contamination that is assumed in the current PWC model. USGS pesticide use estimates are likely a conservative representation of pesticide inputs because they represent only agricultural applications and ignore pesticide applications in homes and industry.

Estimated environmental concentrations from contaminant fate and transport models are favored ways to characterize exposure risk by regulatory agencies because they are low cost, low effort, and provide consistent methodology for estimates across compounds^15^. Currently, these models represent the best methods that have been developed to estimate concentrations of contaminants in the environment. However, these models stand to be improved to increase the accuracy of predictions. We demonstrate that not only are pesticide maximum EECs produced by the PWC model poor characterizations of worst-case exposures, but they also perform poorly at predicting concentrations of pesticides in their intended lentic systems across pesticide types. Estimates of field concentrations in lentic systems can be improved by leveraging large datasets of measured environmental concentration and accounting for sampling effort in validation of models. In addition, including landscape-level contaminant use as a proxy for multiple-sources of contamination can improve PWC model predictions. Scientists active in the development of environmental fate and transport models recognize the importance of including multiple sources of contamination. For instance, models widely used in the United States and Europe incorporate multiple point sources of contamination including the Soil and Water Assessment Tool (SWAT)^31^, ChimERA Fate^32^, and Stream-EU^33^. The inclusion of field survey information and landscape-level use for pesticides is easily accomplished because these data are already included separately in the most current ecological risk assessments used for pesticide regulation^34^. In general, because of environmental laws and regulation requiring reporting of pollution, including the Emergency Planning and Community Right-to-Know Act, the Resource Conservation and Recovery Act, the Toxic Substances Control Act, the Clean Water Act, and the Clean Air Act, there is a clear understanding of the identity and amounts of multiple point sources of many contaminants from industry and agriculture. So, the amounts of contaminants released into the environment at the landscape-level could be feasibly incorporated into EEC models for non-pesticide contaminants as well.

Given our results, the next step for improvement of the PWC model would be for EPA staff members to directly include pesticide use in the mechanistic model. Access to the proprietary computer code that underlies the PWC model prevented us from doing so in the current study. Improving the understanding of the determinants of maximum concentrations of pesticides in lentic systems is not only important for improving exposure characterization as a part of federal ecological risk assessment, but is also critical for the understanding and protecting small freshwater bodies which provide critical habitat to communities of plants and animals^14,35^ and serve an underestimated role in the functioning of ecosystems^36^. Improvement of contaminant fate and distribution models used in federal risk assessments and in the development of regulations is critical if we are to use the best science available to make data driven policy decisions.

## Acknowledgments

We appreciate the advice of D. Young concerning the implementation of EPA’s PWC models, and M. Mahon concerning coding for figures. This work was supported by funds from the National Science Foundation (EF-1241889, EEID-1518681), the National Institutes of Health (R01GM109499, R01TW010286-01), the U.S. Department of Agriculture (NRI 2009-35102-0543).

## Supporting Information

Data used in the current analyses are provided in the supporting information.

